# Determinants of target prioritization and regulatory hierarchy for the bacterial small RNA SgrS

**DOI:** 10.1101/221978

**Authors:** Maksym Bobrovskyy, Jane K. Frandsen, Jichuan Zhang, Anustup Poddar, Muhammad S. Azam, Tina M. Henkin, Taekjip Ha, Carin K. Vanderpool

**Affiliations:** Department of Microbiology, University of Illinois at Urbana-Champaign, 601 S. Goodwin Ave., Urbana, Illinois 61801; Present address: Department of Microbiology, The University of Chicago, 920 E. 58^th^ St., Chicago, Illinois 60637; Department of Microbiology and Center for RNA Biology, The Ohio State University, Columbus, Ohio 43210; Biochemistry Program, The Ohio State University; Department of Biophysics and Biophysical Chemistry, Johns Hopkins University, Baltimore, Maryland, USA 21205; Howard Hughes Medical Institute, Baltimore, Maryland, USA 21205

## Abstract

The mechanisms by which small RNA (sRNA) regulators select and prioritize target mRNAs remain poorly understood, but serve to promote efficient responses to environmental cues and stresses. We sought to uncover mechanisms that establish regulatory hierarchy for a model sRNA, SgrS, found in enteric bacteria and produced under conditions of metabolic stress when sugar transport and metabolism are unbalanced. SgrS post-transcriptionally controls a nine-gene regulon to restore growth and homeostasis under stress conditions. An *in vivo* reporter system was used to quantify SgrS-dependent regulation of target genes and established that SgrS exhibits a clear preference for certain targets, and regulates those targets efficiently even at low SgrS levels. Higher SgrS concentrations are required to regulate other targets. The position of targets in the regulatory hierarchy is not well-correlated with the predicted thermodynamic stability of SgrS-mRNA interactions or the SgrS-mRNA binding affinity as measured *in vitro*. Detailed analyses of SgrS interaction with *asd* mRNA demonstrate that SgrS binds cooperatively to two sites and remodels *asd* mRNA secondary structure. SgrS binding at both sites increases the efficiency of *asd* mRNA regulation compared to mutants that have only a single SgrS binding site. Our results suggest that sRNA selection of target mRNAs and regulatory hierarchy are influenced by several molecular features. The sRNA-mRNA interaction, including the number and position of sRNA binding sites on the mRNA and cofactors like the RNA chaperone Hfq, seem to tune the efficiency of regulation of specific mRNA targets.

**IMPORTANCE:** To survive, bacteria must respond rapidly to stress and simultaneously maintain metabolic homeostasis. The small RNA (sRNA) SgrS mediates the response to stress arising from imbalanced sugar transport and metabolism. To coordinate the stress response, SgrS regulates genes involved in sugar uptake and metabolism. Intrinsic properties of sRNAs such as SgrS allow them to regulate extensive networks of genes. To date, sRNA regulation of targets has largely been studied in the context of “one sRNA-one target”, and little is known about coordination of multi-gene regulons and sRNA regulatory network structure. Here, we explore the molecular basis for regulatory hierarchy in sRNA regulons. Our results reveal a complex interplay of factors that influence the outcome of sRNA regulation. The number and location of sRNA binding sites on mRNA targets and the participation of an RNA chaperone dictate prioritized regulation of targets to promote an efficient response to stress.

## INTRODUCTION

Bacteria live in diverse niches, often encountering rapidly changing and stressful environments. Bacterial stress responses can mitigate the negative effects of stress on cell structure and function. Stress responses are usually coordinated by molecules, either RNAs or proteins, that alter expression of a regulon comprised of multiple genes. Coordinated control of the regulon prepares the cell to survive or adapt to the stress (1). Proteins control expression of target regulons by binding to DNA sequences and modulating the frequency of transcription initiation, whereas RNAs often modulate gene expression post-transcriptionally. A prevalent type of RNA regulator in bacteria is referred to simply as small RNA (sRNA). The sRNAs are often produced in response to a particular stress, and regulate target mRNAs through base pairing interactions that modify mRNA translation or stability (2, 3). Hundreds of sRNAs have been identified in diverse bacteria (4–6). While the majority of sRNAs have not been characterized, many studies suggest that sRNA regulatory networks are as extensive and complex as those controlled by proteins (7, 8).

A large body of work has illuminated base pairing-dependent molecular mechanisms of post-transcriptional regulation by sRNAs (9, 10). The sRNA SgrS (sugar-phosphate stress sRNA) has been an important model for discovery of both negative and positive mechanisms of target mRNA regulation. SgrS is induced in response to metabolic stress associated with disruption of glycolytic flux and intracellular accumulation of sugar phosphates (also referred to as glucose-phosphate stress) (11, 12). SgrS regulates at least 9 genes and promotes recovery from glucose-phosphate stress. SgrS-dependent repression of mRNAs encoding sugar transporters (*ptsG*, *manXYZ*) reduces uptake of sugars to prevent further sugar-phosphate accumulation (Fig. 1) (11, 13, 14). Activation of a sugar phosphatase (*yigL*) mRNA promotes dephosphorylation and efflux of accumulated sugars (15), and repression of other mRNAs is hypothesized to reroute metabolism to promote recovery from stress (Fig. 1) (16). Each target of SgrS is regulated by a distinct molecular mechanism. How different mechanisms of regulation yield effects of variable magnitude with respect to mRNA stability and translation is an open question.

**Figure 1.**
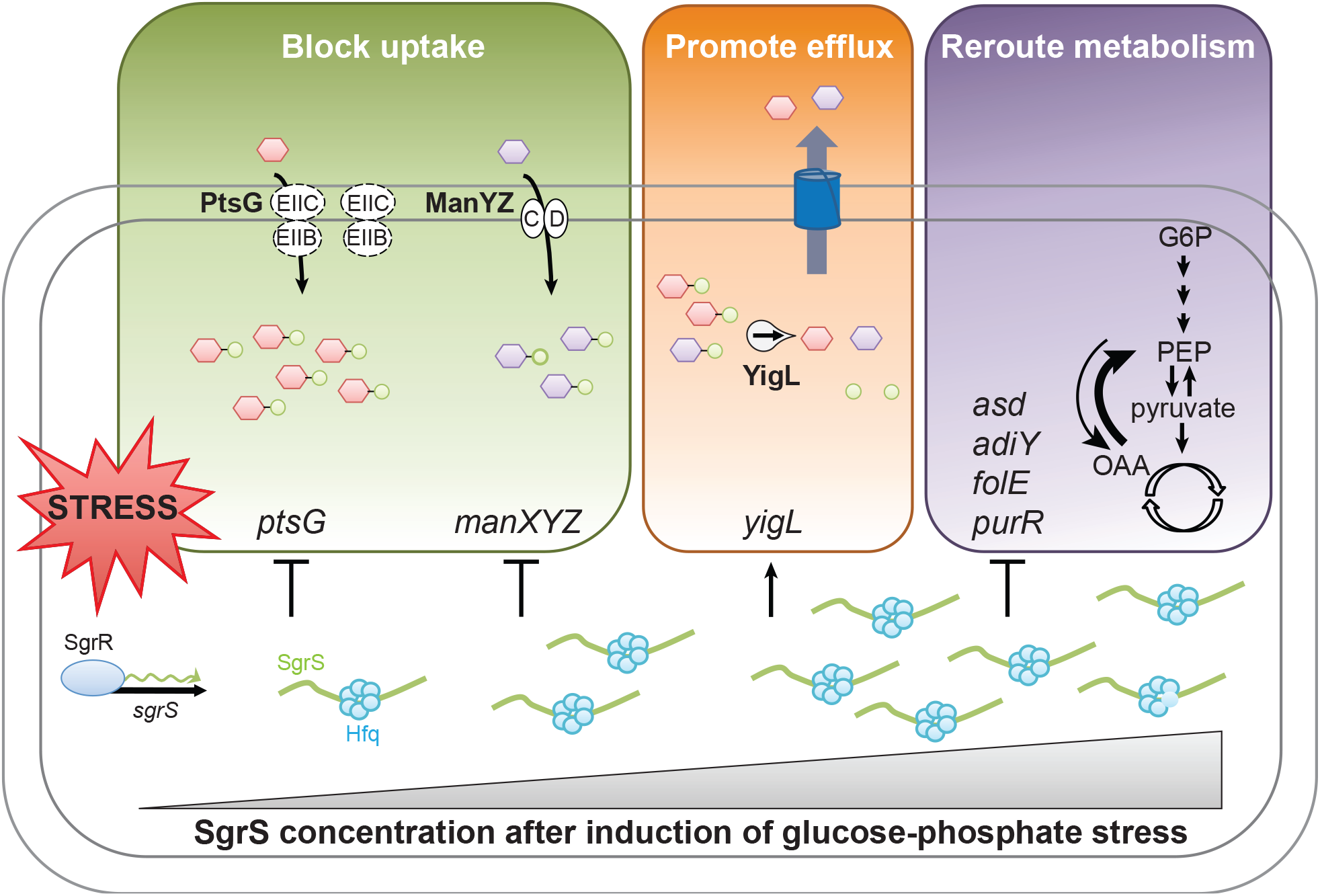
Model for SgrS target prioritization during glucose-phosphate stress. Glucose or the analogs αMG and 2-deoxyglucose are phosphorylated during transport through the phosphotransferase system proteins EIICB^Glc^ (PtsG) or EIICD^Man^ (ManYZ). If sugar-phosphates are not metabolized, the glucose-phosphate stress response is triggered, and the transcription factor SgrR becomes active and promotes *sgrS* transcription. The RNA chaperone Hfq promotes SgrS-mediated translational repression of *ptsG* and *manXYZ* mRNAs, reducing synthesis of sugar transporters. SgrS stabilizes *yigL* mRNA, promoting sugar phosphatase (YigL) synthesis. SgrS-mediated repression of *asd*, *purR*, *folE* and *adiY* likely reroutes metabolism to restore homeostasis during stress recovery. The hypothetical sequence of regulatory events following stress induction is represented from left to right as SgrS levels increase over time. When SgrS concentrations are low, only the highest priority targets are regulated. When stress persists and concentrations of SgrS increase, lower priority targets are regulated.

Temporally-ordered and hierarchical patterns of gene regulation carried out by protein transcription factors have been characterized in many systems (17–20). These regulatory patterns allow cells to respond efficiently to environmental signals by prioritizing induction or repression of products needed to respond to those signals. Protein regulators establish a hierarchy of regulation based on their affinities for binding sites in the operator regions of different target genes. As the concentration of active regulator increases, genes are sequentially regulated based on binding site affinity (21). There is growing evidence that sRNAs also regulate their target genes hierarchically (22, 23). However, the mechanisms involved in establishing and maintaining prioritized regulation of sRNA targets are not known.

We hypothesize that the temporal progression of target regulation by SgrS is optimized to promote efficient recovery from glucose-phosphate stress (Fig. 1). To test this hypothesis, we first defined the efficiency of SgrS regulation of each target and found that SgrS indeed prioritizes regulation of some targets over others. We examined the factors that determine regulatory efficiency, including the arrangement and strength of SgrS target binding sites and the roles of other factors like RNase E and Hfq.

Detailed characterization of a specific SgrS-mRNA target interaction revealed cooperative binding of SgrS to two binding sites and a requirement for both binding sites for maximal SgrS-dependent regulation. Collectively, our results uphold the hypothesis that sRNAs regulate expression of genes in their target regulons hierarchically, and that this is influenced by features of each sRNA-mRNA pair and different molecular mechanisms of regulation that precisely determine the regulatory priority for each target.

## RESULTS

### SgrS differentially regulates targets at the level of translation

Previous studies suggested the possibility of a regulatory hierarchy for the small RNA SgrS, which regulates a diverse set of mRNA targets (10, 11, 13–15). To study this, we used a two-plasmid system to control expression of SgrS and target translational fusions (Fig. 2A). All target transcript fragments fused to *gfp* contain experimentally-confirmed SgrS binding sites. To quantify translational regulation by SgrS and facilitate comparisons of regulatory efficiency among targets, we analyzed the data as described previously (23). Activity of reporter fusions was measured by monitoring GFP fluorescence over time. By plotting the GFP fluorescence (RFU) as a function of growth (OD_600_) for target-gfp fusions in the absence of SgrS, we defined “basal activity” at different inducer concentrations (example in Fig. S1A). This method of quantifying fusion activity accounts for the fact that fluorescence levels are not directly proportional to inducer concentrations ((23) and Fig. S1A). While the absolute values for basal activity differ among different fusions, all fusions responded to induction in a dose-dependent manner (Fig. S2A). Similar plots (RFU/OD_600_) were generated for each fusion in the presence of SgrS (examples Fig. S1B-F). We define “regulated activity” as the slope of the curve (RFU/OD_600_) under conditions where both the fusion and SgrS are induced (example in Fig. S1B). As levels of SgrS increase, clear patterns of repression or induction are seen for all target fusions (Figs S1B-F and S2B-F).

**Figure 2.**
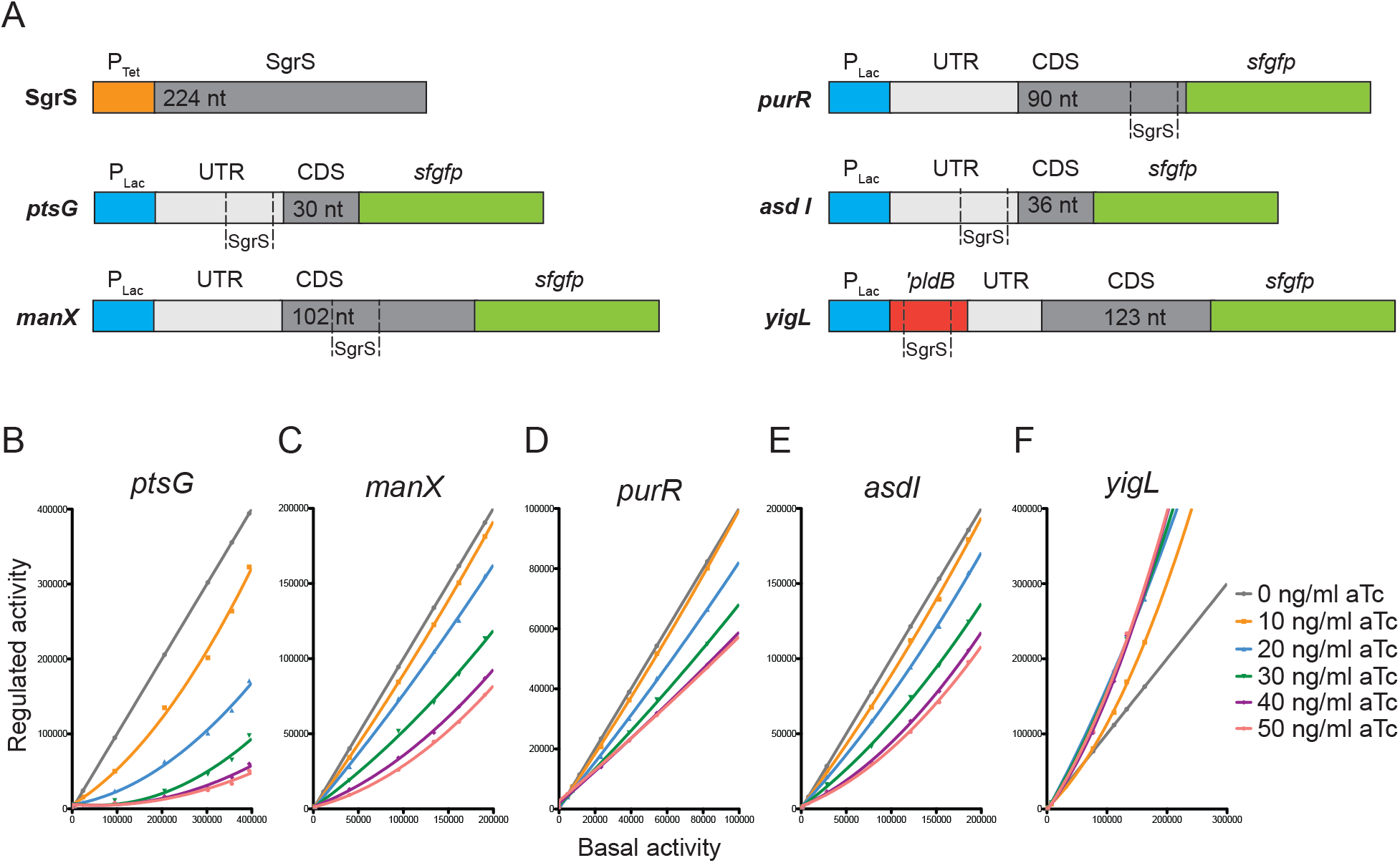
Efficiency of target regulation by SgrS. A) Representation of genetic constructs in two compatible plasmids used to study target regulation by SgrS. One plasmid contains full-length SgrS under the control of the aTc-inducible P_Tet_ promoter. A second plasmid contains a P_Lac_ promoter and the relevant region encoding each SgrS target (including the SgrS binding site) translationally fused to a superfolder *gfp* reporter gene. B-F) Regulated activity was plotted as a function of basal activity (see text for description) for (B) *ptsG*, (C) *manX*, (D) *purR*, (E) *asdI*, and (F) *yigL* fused to *sfgfp* reporter gene. Without SgrS-mediated regulation, we obtained a line with a slope =1. The plots with slopes <1 indicate repression of (B) *ptsG*, (C) *manX*, (D) *purR* and (E) *asdI* by SgrS. The plot with a slope of >1 is indicative of activation of (F) *yigL*.

To define the efficiency of regulation of each target, we plotted regulated activity as a function of basal activity for *ptsG, manX, asdI, purR*, and *yigL*. When there is no SgrS-mediated regulation, a line with a slope of 1 is seen for all targets (Fig. 2B-F). Slopes less than 1 indicate that the fusion is repressed by SgrS. This is true for *ptsG, manX, asdI* and *purR* reporter fusions (Figure 2B-E). Slopes greater than 1 are indicative of activation by SgrS, as seen for *yigL* (Fig. 2F). The magnitude of regulation was responsive to SgrS levels over the range of SgrS inducer (aTc) concentrations (Fig. 2B-E). In contrast, for for *yigL*, the magnitude of activation did not increase beyond a maximal level obtained at 20 ng/mL of inducer (Fig. 2F). While the basis for this difference is unclear, it is likely to reflect the inherently different molecular mechanisms of regulation, *i.e.*, mRNA stabilization for *yigL* and translational repression for other targets.

We then compared regulatory efficiency of targets at different levels of SgrS induction. At the two lowest levels of SgrS induction (10ng/mL and 20 ng/mL aTc), only *ptsG* and *yigL* showed substantial repression and activation, respectively (Fig. 3A, B). In contrast, *manX, asdI* and *purR* fusions yielded curves whose slopes remained at ~1, indicating no regulation at lower levels of SgrS. Our interpretation is that *ptsG* and *yigL* are high-priority targets of SgrS, since they are regulated preferentially when SgrS levels are low. With increasing SgrS levels (20-50 ng/ml aTc), regulation of “weaker” targets *manX, asdI* and *purR* became apparent (Fig. 3C, D, E). As SgrS levels increased, *ptsG* repression became more efficient up to a maximal repression at 40 ng/mL of aTc, and it remained the most strongly repressed target at all levels of SgrS. Collectively, these data suggest that SgrS targets are preferentially regulated in the following order: 1/2) *ptsG* and *yigL*, 3) *manX*, 4) *asdI*, and 5) *purR* (Fig. 3A-E). We hypothesize that the position of each target within the regulatory hierarchy is determined by characteristics of SgrS-target mRNA interactions and the mechanism of SgrS-dependent regulation.

**Figure 3.**
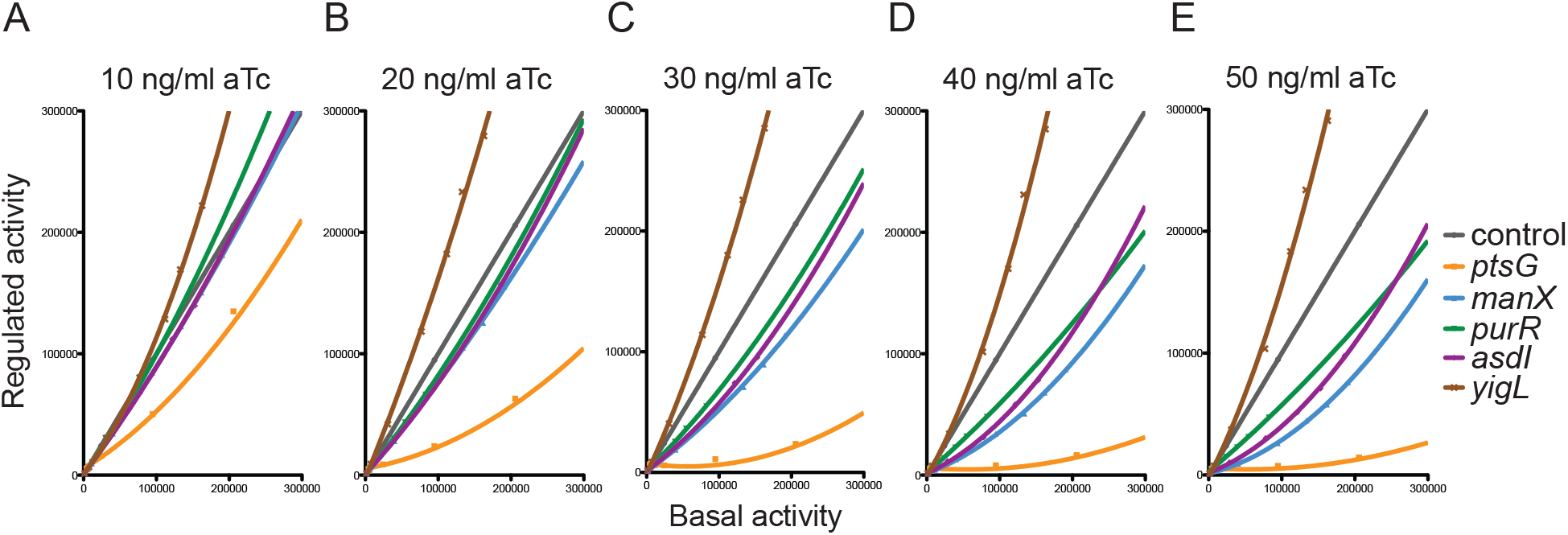
Regulatory hierarchy established by SgrS. Regulated activity was plotted as a function of basal activity for *ptsG, manX, purR, asdI*, and *yigL* fusions. Lack of SgrS regulation is indicated by a line with a slope =1. The plots with slopes <1 indicate repression (*ptsG*, *manX*, *purR* and *asdI*) by SgrS. The plot with slope >1 indicates activation (*yigL)*. Target fusion activity was monitored at different levels of SgrS induction by aTc: (A) 10 ng/ml, (B) 20 ng/ml, (C) 30 ng/ml, (D) 40 ng/ml, (E) 50 ng/ml.

### Differences in binding affinity *in vitro are* not correlated with regulation efficiency

One of the initial steps in sRNA-mediated regulation is formation of base-pairing interactions with the target mRNA. Binding of the sRNA with its target mRNA is dependent on sequence complementarity and RNA secondary structure. We examined the characteristics of SgrS-target mRNA binding *in vitro* to determine whether the strength of binding is correlated with the target hierarchy.

Electrophoretic mobility shift assays (EMSAs) were performed to measure binding of SgrS to its target mRNAs *ptsG, manX, purR, yigL* and *asd*. SgrS bound to *ptsG* with a *K*_D_ of 0.11 ± 0.01 μM (Fig. 4A, B), which was lower than *K*_D_s for SgrS binding to most of the other targets (Fig. 4A-E). SgrS-manX mRNA binding had a *K*_D_ of 19.7 ± 2.78 μM (Fig. 4A, C), which is weaker than the interaction with *ptsG* (Fig. 4B), but stronger compared to *purR* and *yigL* (Fig. 4A). Three different fragments of *asd* mRNA were tested, because previous work demonstrated that SgrS pairs at two distinct sites on *asd* mRNA (16). The first site, *asdI*, is adjacent to the ribosome binding site and is sufficient for modest SgrS-dependent translational repression. The second site, *asdII*, is in the coding sequence of *asd*, 60-nt downstream of the start codon. When both sites are present (RNA *asdI-II)*, stronger SgrS-dependent translational repression was observed (16). Surprisingly, while *asdI* RNA (containing only the upstream SgrS binding site) regulation was less efficient compared to that of *manX* (Fig. 3A-E), *in vitro* it bound SgrS more strongly with a *K*_D_ of 0.15 ± 0.04 μM (Fig. 4A, D), which is comparable to SgrS-ptsG binding (Fig. 4A, B). SgrS interaction with *asdII* was very weak (Fig. 4A). We could not determine *K*_D_ values for SgrS interaction with *asdII, purR* and *yigL*, due to limitations in obtaining high enough concentrations of RNA, but it is apparent that SgrS binding to these targets is much weaker compared to *ptsG, manX* and *asdI* (Fig. 4A).

**Figure 4.**
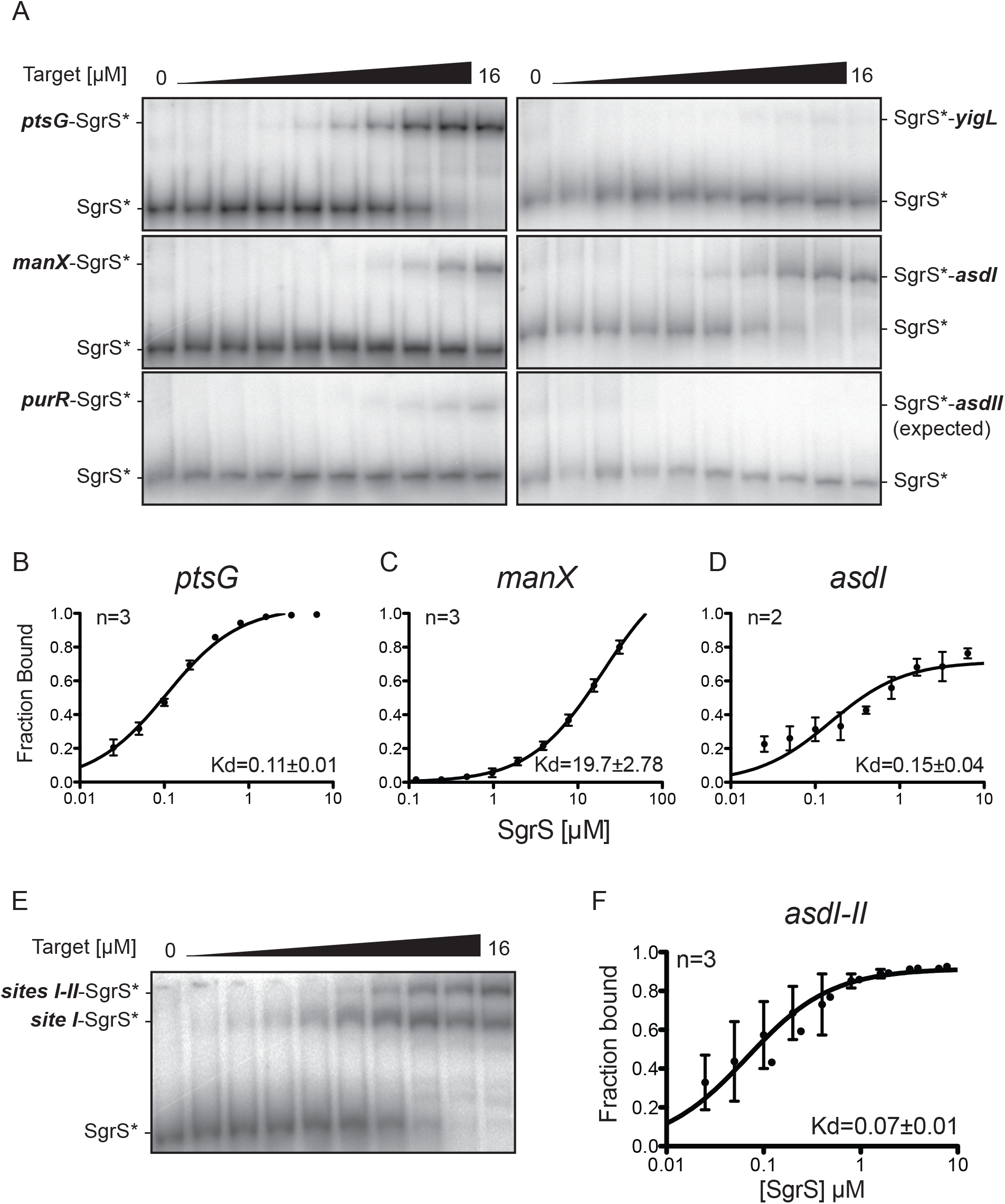
SgrS binding to target mRNAs *in vitro*. A) SgrS was labeled with ^32^P and incubated with unlabeled target transcripts at final concentrations of 0 μM - 16 μM. Electrophoretic mobility shift assays (EMSAs) were performed after incubating full-length SgrS (+1 to +227) with its target transcripts (A) *ptsG* (+1 to +240), *manX* (+1 to +240), *purR* (+1 to +230), *yigL* (-191 to +50 relative to ATG translation start of *yigL*), *asdI* (+1 to +110), and *asdII* (+71 to +310). B-D) Target transcripts (B) *ptsG* (+1 to +240), (C) *manX* (+1 to +240), (D) *asdI* (+1 to +110) were labeled with ^32^P and incubated with unlabeled SgrS. EMSAs were performed to resolve complex formation. Band densities were measured for replicate experiments (n, top left) and plotted to determine dissociation constant (*K*_D_, bottom right) values for (B) *ptsG*, (C) *manX*, and (D) *asdI*. E) EMSA of radiolabeled SgrS in the presence of increasing concentrations of *asdI-II* transcript. Shift in mobility corresponding to one or two SgrS bound to *asdI-II* is denoted as Site I-SgrS* and Sites I-II-SgrS* respectively. F) Quantification of SgrS binding with radiolabeled *asdI-II* (+1 to +240), as described above.

Results of EMSAs with SgrS and *asdI-II* (containing both SgrS binding sites) revealed apparent binding cooperativity. SgrS binding to *asdI-II* has a *K*_D_ of 0.07 ± 0.01 μM (Fig. 4E, F), slightly lower than that of SgrS-ptsG mRNA binding. Moreover, we observed two shifted species that correspond to binding of either one or two SgrS sRNAs to a single *asdI-II* RNA (Fig. 4E).

### Structural analyses of *SgrS-asd* mRNA interactions

Our data thus far indicate that SgrS regulates mRNA targets in a hierarchical fashion (Figs. 2, 3). However, SgrS-mRNA binding affinities do not explain the target hierarchy, as SgrS-ptsG mRNA and SgrS-asd mRNA interactions have very similar *K*_D_s, but *ptsG* is much more efficiently regulated than *asd* at all concentrations of SgrS (Fig. 3). To further understand the features that influence the efficiency of target regulation, we performed more detailed analyses of SgrS-asd mRNA interactions.

Previous work demonstrated that SgrS binding site I encompasses nt +31 to +49 and site II encompasses nt +110 to +127 ((16), Fig. 5A). We used IntaRNA (24, 25) to predict the free energy (ΔG) for SgrS interactions with *asd* mRNA segments containing both sites, or each site individually (Fig. 5B). IntaRNA accounts for the energy of opening double-stranded RNA secondary structure, to make sequences accessible for pairing. We first analyzed SgrS interactions with *asdI-II* mRNA (nt +1 to +240), which we denote as “structured.” Interaction of SgrS with *asd* site I has a predicted ΔG of −10.5 kcal/mol, while SgrS pairing with site II has a predicted ΔG of −1.1 kcal/mol (Fig. 5B, structured). The ΔG for interactions between SgrS and the isolated binding sites are −18 kcal/mol for site I and −7.4 kcal/mol for site II (Fig. 5B, isolated). These predictions suggest that SgrS interaction with site II is less favorable, particularly in the context of the longer structured *asd* mRNA.

**Figure 5.**
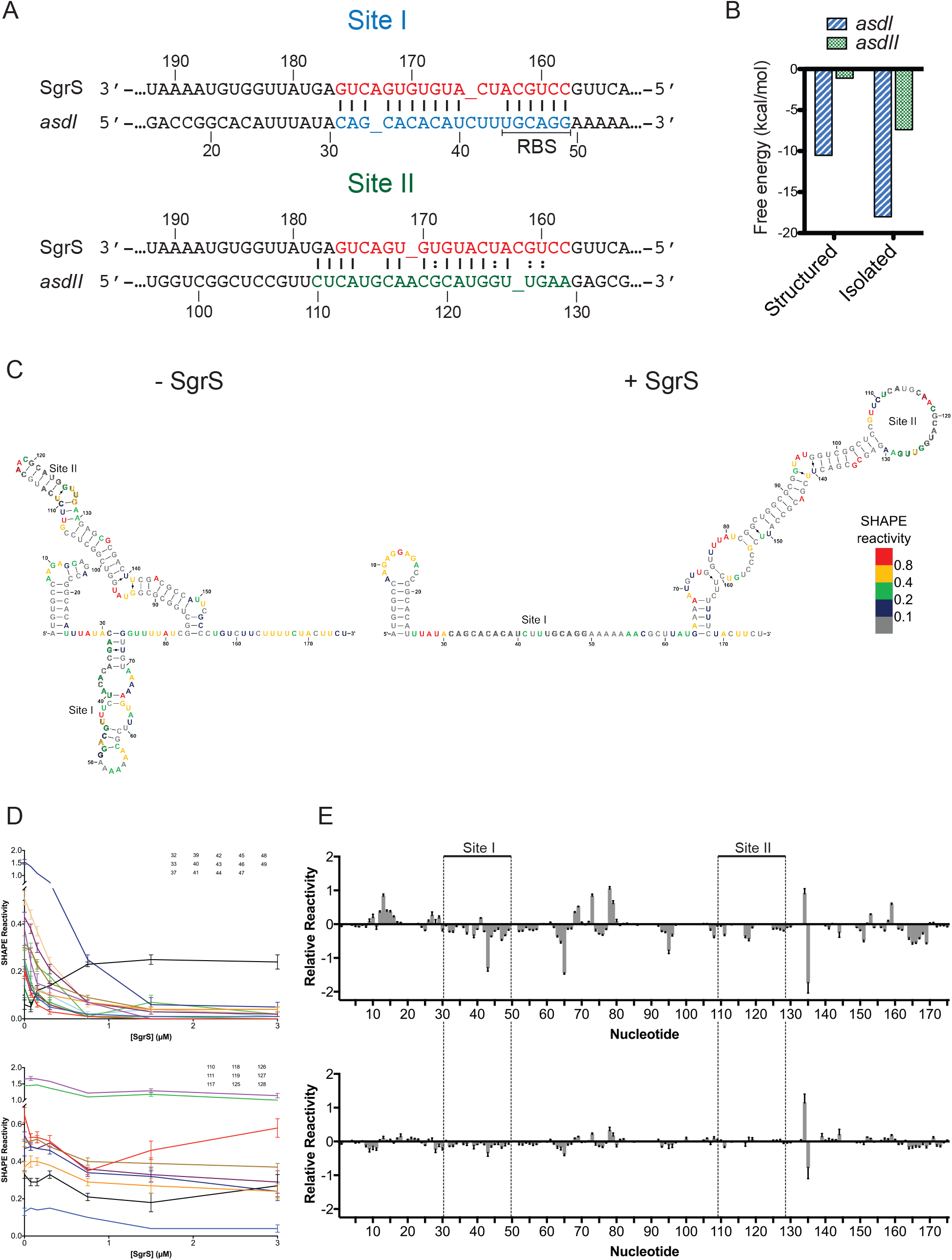
Secondary structure of 5’ end of *asd*. A) Diagram showing base-pairing interactions of SgrS with binding sites I and II of *asd* mRNA. B) Energy of interaction predicted by IntaRNA (26). “Structured” indicates pairing between full length SgrS (+1 to+227) and *asdI-II* (+1 to +180). Plotted is the energy of interactions at either site I (*asdI*)or site II (*asdII*). “Isolated” indicates interactions between isolated binding sites: SgrS (+158 to +176) with *asdI* (+31 to +49) and SgrS (+158 to +178) pairing with *asdII* (+110 to +129). C) The structure of the *asdI-II* RNA alone or in complex with SgrS was probed with NMIA and the modified RNA was analyzed by primer extension inhibition. SHAPE reactivity (difference between the frequency of primer extension products at each nucleotide in +NMIA vs. -NMIA samples) was then used as a parameter in the Vienna RNAprobing WebServer (47) to predict the secondary structure of the *asdI-II* RNA. Colors indicate SHAPE reactivity as follows: red, highly reactive (≥ 0.8); gold, reactive (0.4-0.79); green, moderately reactive (0.2-0.39); blue, minimally reactive (0.1-0.19); grey, unreactive (< 0.01). Distinct structures were observed in the absence of SgrS and in the presence of saturating concentrations of SgrS. The SHAPE reactivity of *asdI-II* RNA alone (left) or in the presence of 5-fold excess SgrS (right) is mapped to the predicted secondary structures. (D) SHAPE reactivity as a function of SgrS concentration for each binding site (top, site I; bottom, site II). Only nucleotides with a significant (≥ 0.1) change in reactivity are shown. Error bars denote SEM, n = 9. (E) Relative SHAPE reactivity (difference in the SHAPE reactivity in the presence of SgrS vs. the absence of SgrS) of the *asdI-II* RNA in the presence of wild-type (top) or mutant (bottom) SgrS. Error bars denote SEM, n = 9 (WT), 6 (MT). The *asdI-II* RNA nucleotides are numbered below the X-axis and the SgrS binding sites are indicated.

We investigated the structure of *asdI-II* with selective 2’-hydroxyl acylation analyzed by primer extension (SHAPE), in which flexible nts are modified by N-methylisotoic anhydride (NMIA), while nts constrained in helices are not reactive. In the absence of SgrS, the sequence encompassing the *asd* ribosome binding site (+44 to +50) is located within a structured loop (+36 to +69) on top of a short stem (+31 to +35 pairing with +70 to +74) (Fig. 5C, Fig. S3). The nts in site I (+31 to +49, Fig. 5A) are located on the 5’ side of the stem-loop structure. Most of the nts in this structure are reactive, which is indicative of a flexible conformation that is accessible for ribosome binding or base pairing with the seed sequence of SgrS (Fig. 5C). The seed interaction of SgrS is likely to promote opening of the structure. Downstream of the site I stem-loop structure is a highly structured second stem (+83 to +155) that contains site II in the apical region (+110 to +129) (Fig. 5C, Fig. S3). Site II is sequestered in a helix and would not be accessible to base pair with SgrS (Fig. 5C). In light of binding cooperativity observed in Fig. 4E, we hypothesize that SgrS pairing with site I induces rearrangement of *asd* mRNA secondary structure to facilitate interaction with site II.

We next used SHAPE to probe changes in the *asdI-II* structure in the presence of SgrS. The reactivity of site I nt +31 to +49 decreased as the concentration of SgrS increased (Fig. 5D), with the exception of nt +41 which is not predicted to base pair with SgrS (Fig. 5A). The SHAPE reactivity plateaued between 5- and 10-fold excess SgrS (Fig. S3E,F), which suggests that binding to site I was saturated. This is consistent with a strong base-paring interaction between SgrS and site I. In contrast, the reactivity of the site II nts +110 to +129 decreased more slowly and to a lesser extent (Fig. 5D), consistent with a weaker and cooperative interaction. Fewer site II nts showed changes in SHAPE reactivity upon addition of SgrS; this is likely to be due to the highly structured nature of site II in the absence of SgrS.

The reactivity of nts outside of the SgrS binding sites also changed in the presence of SgrS (Fig. 5E). In contrast, when a mutant SgrS that is not predicted to bind to *asdI-II* was used, minimal changes in SHAPE reactivity were observed, which suggests that the changes in the presence of wild-type SgrS are due to the specific interactions between SgrS and *asdI-II* mRNA and not to the presence of additional RNA in the system (Fig. 5E). These results indicate that SgrS binding changes the overall structure of the *asd* RNA. A secondary structure predicted using the SHAPE data suggests that these changes are limited to opening the SgrS binding sites and extending the site II stem (Fig. 5C). We note an important caveat to these analyses. The structure prediction algorithms were not designed to account for intermolecular interactions, so this analysis may not be able to capture the *in vivo* relevant structure of *asd* mRNA in complex with SgrS. Nonetheless, SHAPE data are consistent with other analyses in demonstrating binding of SgrS to *asd* mRNA, prominently at site I and to a lesser extent at site II.

### Optimal repression by SgrS involves both pairing sites within *asd* mRNA

To further investigate the role of the two SgrS pairing sites on *asd*, we performed stochastic optical reconstruction microscopy (STORM) coupled with single-molecule *in situ* hybridization (smFISH) to monitor SgrS regulation of *asd-lacZ* variants *asdI, asdII*, and *asdI-II* (Fig. 6A) at single molecule resolution. In these experiments, bacteria were grown in the presence of L-arabinose to induce expression of chromosomal *asd-lacZ* variants, and glucose-phosphate stress was induced for 10 min by the addition of 1% a-methyl D-glucopyranoside (αMG). 3D super-resolution images show *asd-lacZ* mRNAs (Fig. 6B-D, green) and SgrS (Fig. 6B-D, red), as projected on 2D planes, with cells outlined. Numbers of *asd-lacZ* mRNAs and SgrS sRNAs were counted and represented as “copy number per cell” in histograms, with average copy number per cell indicated above the histogram (Fig. 6B-D). SgrS induction reduced the copy number of *asdI-lacZ* RNA by 3-fold (Fig. 6B, green) when SgrS was induced to high levels after 10 min treatment with αMG (Fig. 6B, red and S4A, B). In contrast, the copy number of *asdII-lacZ* RNAs (Fig. 6C, green) was not strongly affected in the presence of high SgrS levels after αMG treatment (Fig. 6C, red and Fig. S4C, D). Copy numbers of *asdI-II-lacZ* RNA (Fig. 6D, green) were reduced by ~8-fold after 10 min of αMG induction (Fig. 6D, red, Fig. S6E,F). These data demonstrate that both binding sites on *asd* mRNA are important for efficient SgrS-dependent regulation of mRNA stability.

**Figure 6.**
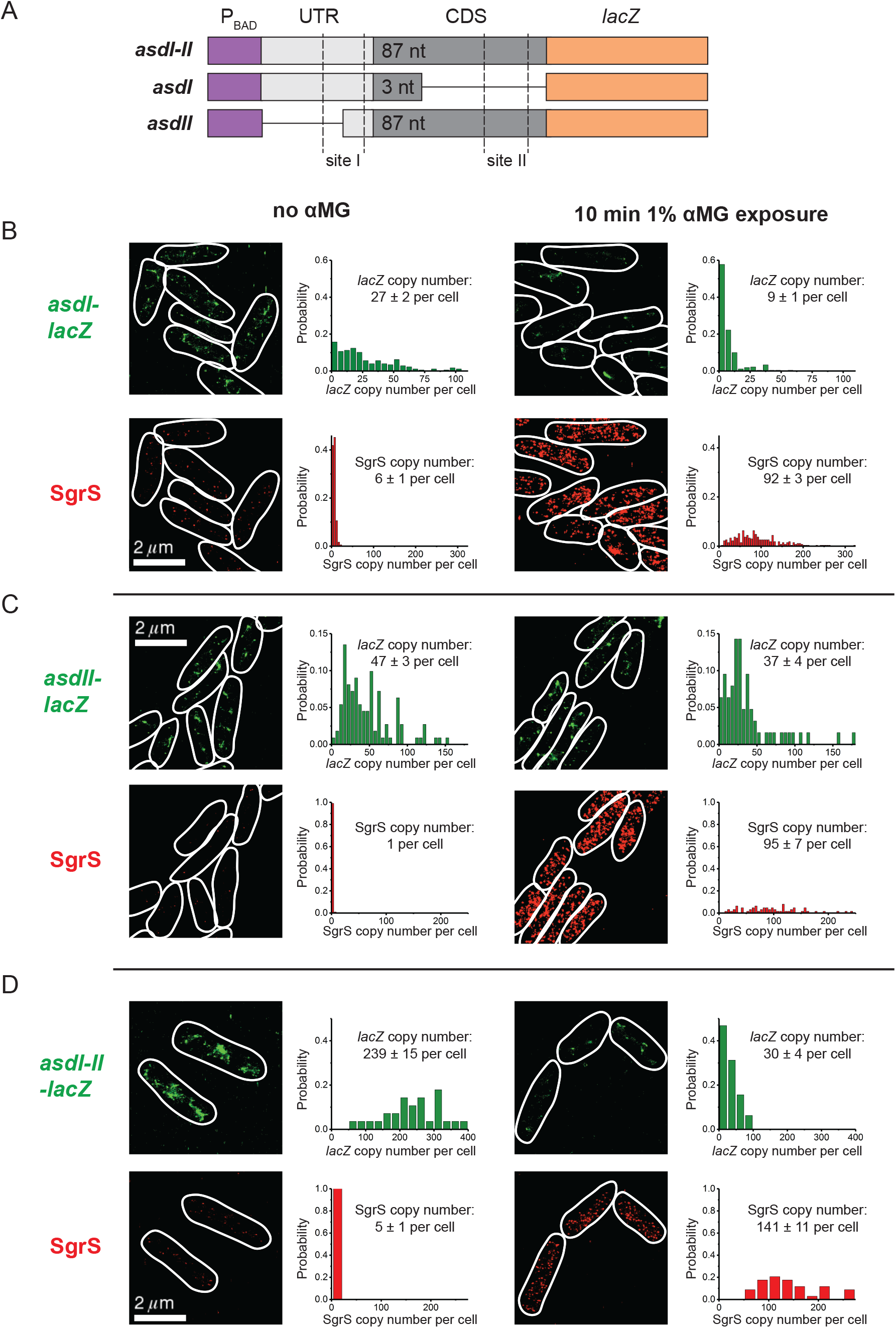
STORM imaging of SgrS regulation of *asd* variants. A) Illustration of *asdI-II*, *asdI* and *asdII* translationally fused to *lacZ* reporter with SgrS binding sites I and II marked. B-D) 2D projection of 3D super-resolution images of SgrS and *lacZ* mRNA for the different *asd-lacZ* variants, labeled by smFISH, before and after 10 min induction with 1% αMG. (B), (C) and (D) correspond to *asdI-lacZ, asdII-lacZ*, and *asdI-II-lacZ* shown in (A). Probability distributions of RNA copy numbers in individual cells for 30-250 cells are plotted next to the representative images.

We next examined the roles of the two SgrS binding sites in the efficiency of translational regulation. SgrS regulation of an *asdI-II* translational fusion was compared to regulation of an *asdI* fusion (Fig. 7A). By plotting regulated activity as a function of basal activity as described above, we determined that SgrS repression of *asdI-II* was more efficient than repression of *asdI* across a range of SgrS expression levels (Fig. 7B), a result in line with our previous study (16). Comparison to other targets indicated that *asdI-II* is regulated more efficiently than *manX, asdI* and *purR*, at all concentrations of SgrS (Fig. 7C).

**Figure 7.**
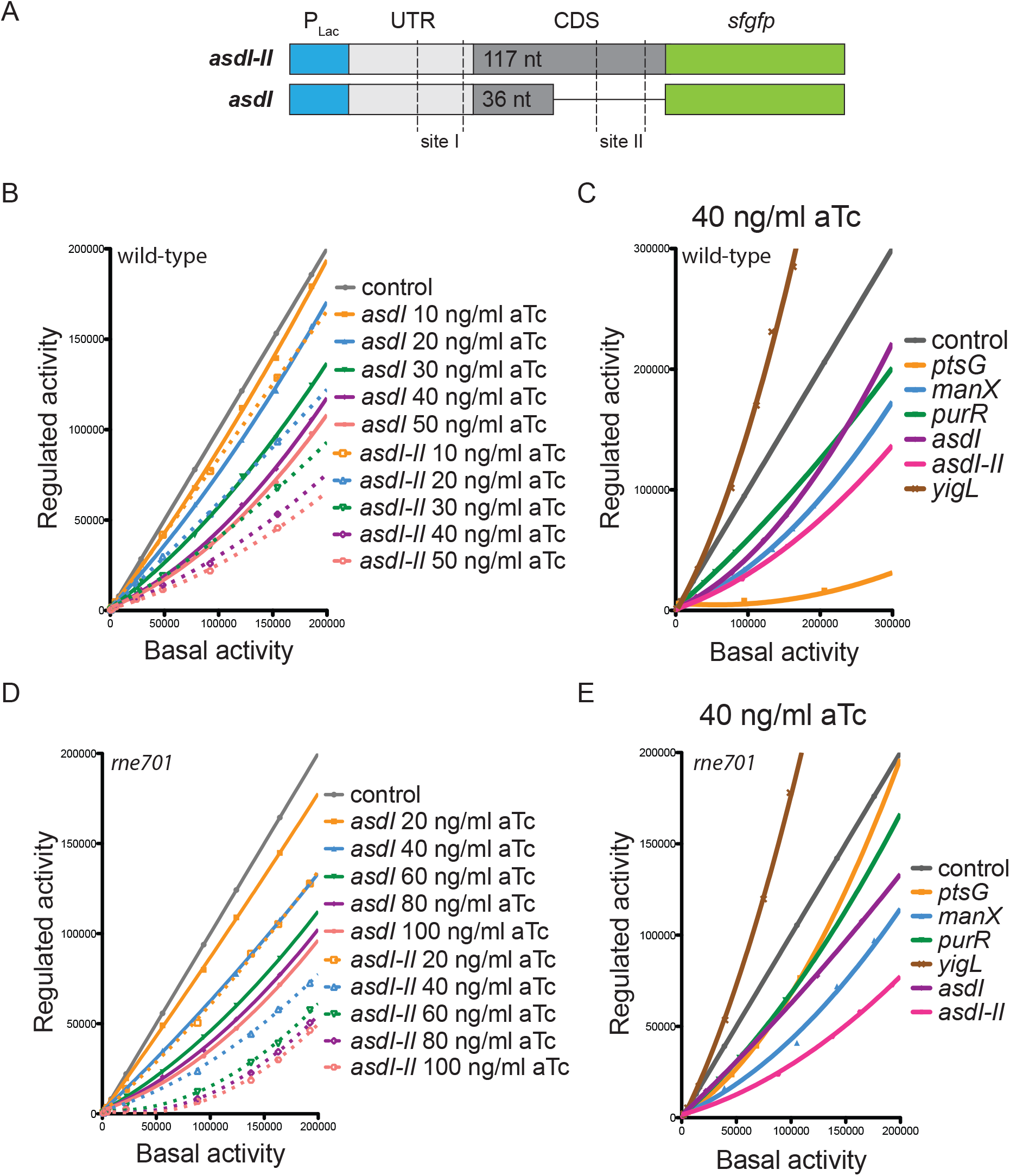
SgrS binding cooperativity allows for improved repression of *asd* translation. A) Illustration of *asdI* and *asdI-II* constructs with SgrS binding sites marked. Graphs show comparison of SgrS regulation of *asdI* and *asdI-II* variants in (B) wild-type and (D) *rne701* mutant by plotting regulated activity over basal activity at various SgrS expression levels (20-100 ng/ml aTc). Regulatory hierarchy of SgrS targets in (C) wild-type and (E) *rne701* mutant strains. Regulation of target genes at one SgrS expression level (40 ng/ml aTc) are compared by plotting regulated activity over basal activity of the *gfp* reporter.

We then compared SgrS regulation of *asdI* and *asdI-II* in the *rne701* mutant strain deficient in degradosome assembly (26). We determined basal activity (Fig. S5A) and regulated activity (Fig. S5B-F) of *asdI* and *asdI-II* translational *gfp* fusions at different levels of SgrS induction. Similar to our data for the wild-type strain (Fig. 7A), SgrS regulated *asdI-II*more efficiently compared to *asdI* in the *rne701* mutant (Fig. 7D). Moreover, when compared to SgrS regulation of other targets, *asdI-II* was repressed most efficiently (Fig. 7E) in the *rne701* mutant. Taken together the data indicate that the second binding site on *asd* mRNA enhances the stringency of SgrS-mediated regulation. Moreover, addition of the second binding site on *asd* changes its regulatory priority relative to other targets in the SgrS regulon.

### Transcriptional regulation of *asd* by SgrS

We observed that the *asdI-II* RNA is more abundant than *asdI* (Fig. 6B, D). Since the constructs used in that experiment were expressed from a heterologous promoter, we postulated that increased levels of *asdI-II* compared to *asdI* mRNA must be due to increased mRNA stability or transcription elongation of the *asdI-II* construct compared to *asdI*. We therefore constructed *asdI* and *asdI-II* transcriptional fusions to *lacZ* expressed from an inducible promoter (Fig. 8A) to test whether SgrS can regulate *asd* transcripts independent of translational regulation. Consistent with observations from smFISH, the *asdI-II-lacZ* transcriptional fusion had substantially higher activity compared to *asdI-lacZ* (Fig. 8B). Repression of *asdI-II* by SgrS was more efficient (3.3-fold repression) than was observed for *asdI* (2.1-fold repression) (Fig. 8B). SgrS still regulated both fusions in the *rne701* mutant strain (Fig. 8B). Importantly, SgrS-dependent degradation of other SgrS targets (*ptsG* mRNA (27) and *manXYZ* mRNA (13, 14)) was abolished in degradosome mutants. Together with our previous work, these observations suggest that SgrS regulates *asd* by two independent mechanisms, translational repression by pairing at site I (directly occluding the ribosome binding site) and reducing mRNA levels by promoting mRNA turnover and/or inhibiting transcription elongation.

**Figure 8.**
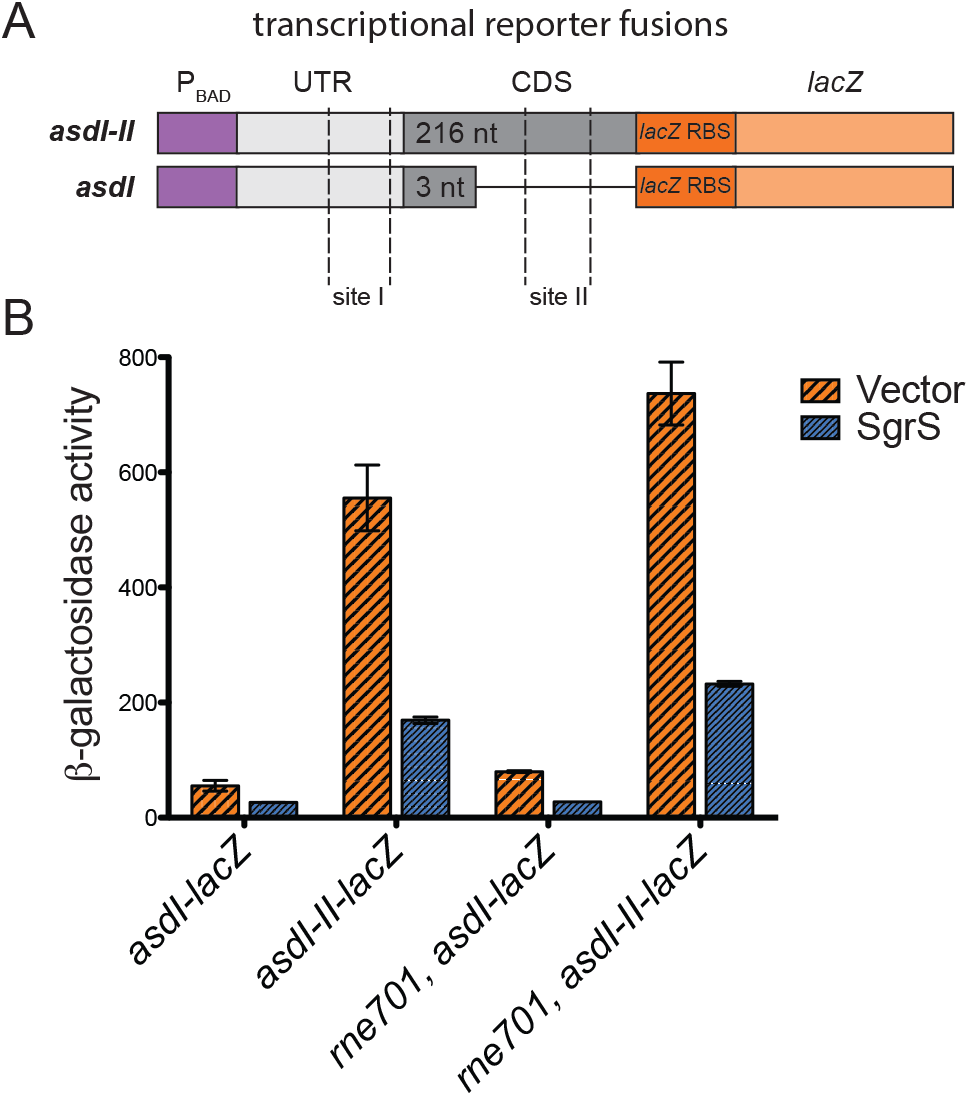
SgrS regulation of transcriptional *asd-lacZ* fusions. β-galactosidase activity of (A) *asdI-lacZ* (+1 to +64) and *asdI-II-lacZ* (+1 to +277) was (B) assayed in response to SgrS expression from a plasmid (and vector control) in WT and *rne701*background strains.

## DISCUSSION

In this study, we set out to define the hierarchy of regulation by a model bacterial sRNA. SgrS is a good model for this study because it has a modestly-sized regulon, and the mechanisms of regulation of several targets have been characterized in detail (13, 15, 16, 28). Our results demonstrate a clear pattern of prioritized regulation of mRNA targets (Fig. 2B-F, Fig. 3A-E). Two targets in particular, *ptsG* and *yigL*, were “high-priority” targets that were efficiently regulated even at low levels of SgrS. Other targets, *manX, purR*, and *asd*, were less impacted by SgrS and were regulated only when SgrS was produced at higher levels.

We investigated features of sRNA-mRNA interactions that could impact the overall efficiency of SgrS-mediated regulation. SgrS-mRNA interactions *in vitro*, as measured by EMSA defined *K*_D_s, were not well-correlated with *in vivo* regulatory efficiency (Fig. 4A-F, Fig. 3A-E). Two targets stood out in the comparison of *in vivo* regulation and *in vitro* SgrS-mRNA interactions. The *yigL* mRNA interaction with SgrS was barely detectable *in vitro* (Fig. 4A), but *in vivo, yigL* translation was maximally activated at low SgrS levels (Fig. 2F). Conversely, the translation of *asdI* was modestly regulated by SgrS *in vivo* (Fig. 2E), but the *in vitro SgrS-asdI* interaction was comparable to that of SgrS-ptsG, which had the strongest *in vivo* regulatory effect.

These apparent contradictions between *in vitro* interactions and *in vivo* regulatory efficiency led us to further explore SgrS regulation of *asd*.

Previous work demonstrated that SgrS has two binding sites on *asd* mRNA: site I overlaps that *asd* ribosome binding site and site II is ~60 nt downstream in the *asd* coding sequence ((16) and Fig. 5A). EMSAs demonstrated SgrS pairing at site I alone, but pairing at site II alone was undetectable. Binding of SgrS to an *asd* mRNA containing both sites I and II was cooperative (Fig. 4E,F). Structural analyses of *asd*mRNA in the absence and presence of SgrS demonstrated that SgrS indeed pairs preferentially at site I over site II and induces substantial structural rearrangement in the mRNA (Fig. 5C-E, Fig. S3). Quantification of SgrS-dependent degradation of *asd* mRNA showed that site I is important, but sites I and II together promote the most efficient regulation (Fig. 6B-D, Fig. S4). Similar to binding and regulation of mRNA degradation, SgrS interactions at both sites I and II on *asd* mRNA improve the efficiency of translational regulation (Fig. 7B,C, Fig. S2). These results suggest that the number and position of sRNA binding sites on mRNA targets control regulation *in vivo* in ways that could not be predicted based on *in vitro* characterization of sRNA-mRNA binding.

In many cases, sRNA-mediated regulation of translation is thought to indirectly affect mRNA stability by making untranslated mRNA more susceptible to degradation by RNase E. There are also examples of sRNA regulation, including SgrS regulation of *yigL* (15), where modulation of mRNA stability is translation-independent. Truncation of RNase E (encoded by rne), removing the C-terminal scaffold for degradosome assembly, often prevents sRNA-mediated degradation of mRNA targets (14, 29, 30). If translational regulation is the primary function of an sRNA on a given mRNA target, the regulation should be preserved in *rne* mutant backgrounds. For SgrS targets, the regulatory hierarchy is mostly preserved in an *rne701* degradosome mutant background (Fig. 7, compare C and E), suggesting that for most targets, regulation of RNA stability is not the primary mechanism of control by SgrS. Interestingly, the high-priority target *ptsG* was a notable exception. In the wild-type background, *ptsG* is the most efficiently-regulated target (Fig. 7C), whereas in the *rne701* host, it is weakly regulated. This defect could be overcome by increasing SgrS expression levels (Fig. S6B). This result suggests that RNase E-dependent degradation of *ptsG* mRNA is more important for its efficient regulation by SgrS compared to other targets, where efficient regulation does not depend on subsequent target degradation. This is consistent with the fact that *ptsG* mRNA levels decrease at least 10-fold whereas other targets exhibit a modest 2-fold decrease in mRNA levels upon SgrS expression (16). Our recent study quantifying SgrS-dependent mRNA degradation at single molecule resolution indicated that *ptsG* mRNA exhibits faster degradation kinetics than *manXYZ* mRNA (29), which could enhance the efficiency of regulation in a wild-type but not *rne701* mutant background where translational regulation and mRNA degradation are uncoupled.

One of our ultimate goals is to define at a molecular level the mechanisms by which sRNAs select and prioritize regulation of their targets. The current study implicates features of sRNA-mRNA interactions such as number and strength of sRNA binding sites on each mRNA target and accessory factors such as RNase E in dictating regulatory hierarchy. Another factor that is likely to play an important role in setting regulatory priority is the RNA chaperone Hfq. EMSAs demonstrated Hfq binding to *ptsG, manX, purR, yigL, asdI, asdII* and *asdI-II* mRNAs (Fig. S7A) with similar *K*_D_ values for all targets (Fig. S7B). Previous work has shown that sRNAs compete for binding to Hfq, and this competition affects their regulatory ability (31, 32). Different sRNAs can bind to distinct sites on Hfq and this impacts their regulation of particular targets (32, 33). Additional work will be required to determine what role Hfq plays in establishing the hierarchy of regulatory effects in sRNA regulons.

Most sRNA-mRNA interactions are conceived of as single binding site interactions, but we have already identified two SgrS targets that deviate from this model and have shown that additional binding sites can play important roles in regulation and change regulation efficiency (13, 16). We have not yet discovered the specific mechanism of regulation of *asd* mRNA by SgrS, but have shown definitively that both binding sites are required for strong regulation. SgrS-dependent control of both transcriptional and translational *asd* reporter fusions is not impacted in RNase E degradosome deficient strains (Fig. 7B,D, Fig. 8B), suggesting that the regulation is not strictly dependent on translation or mRNA turnover. Future work will test the hypothesis that SgrS acts on *asd* mRNA at the level of transcription elongation, perhaps by an attenuation mechanism.

In *Vibrio*, quorum sensing-regulated Qrr sRNAs regulate multiple targets by distinct mechanisms and differences in those mechanisms influence the dynamics and strength of regulation (22). Strong and rapid regulation is achieved by sRNAs acting catalytically, whereby the sRNA pairs with and promotes mRNA degradation but is then recycled for use on another mRNA target. A sequestration mechanism, where formation of the sRNA-mRNA complex is the terminal outcome of regulation, results in slower and weaker sRNA-dependent regulation of the target mRNA. For Qrr sRNAs, these regulatory mechanisms seem to depend on which region of the sRNA is pairing with a given target and whether the sRNA-mRNA interaction is strong or weak (22). While some of the same features of SgrS-mRNA interactions may be relevant in determining regulatory efficiency, we note that the SgrS seed sequence responsible for pairing with all mRNA targets characterized thus far is encompassed by a short (~20 nt) mostly single-stranded region of SgrS (11, 14–16). Moreover, we did not see a good correlation between strong versus weak binding *in vitro* and *in vivo* regulatory efficiency. It is possible that the “rules” governing regulatory efficiency and specific outcomes are different for each individual sRNA. Detailed analysis of more model sRNAs will be needed to illuminate broad general principles.

Beyond defining interesting molecular features of sRNA-mRNA interactions, defining regulatory hierarchy for sRNA regulons is important for understanding bacterial physiology. The vast majority of sRNA regulons remain undefined, and thus many sRNA functions are unknown. For novel sRNAs, distinguishing high-priority from weaker targets may provide crucial clues to the predominant role of the sRNA in cell physiology. For SgrS, the regulatory hierarchy we have defined here is perfectly consistent with growth studies that demonstrate the primary importance of SgrS regulation of sugar transport and efflux under glucose-phosphate stress conditions (34). The hierarchy of regulation by sRNAs is likely to have evolved to promote rapid and efficient responses to environmental signals that would provide cells with a competitive growth advantage in their specific niche. It will be crucial to develop tools to more rapidly characterize sRNA regulatory hierarchy to better understand the functions of the hundreds of uncharacterized sRNAs in diverse bacteria.

## MATERIALS AND METHODS

### Strain and plasmid construction

Strains and plasmids used in this study are listed in Table S1. All strains used in this study are derivatives of *E. coli* K-12 strain MG1655. Oligonucleotide primers and 5’ biotinylated probes used in this study are listed in Table S2 and were purchased from Integrated DNA Technologies. Chromosomal alleles were moved between strains by P1 *vir* transduction (35) and inserted using λ Red recombination (36, 37).

Translational reporter fusion alleles P_BAD_-*asdI-II-lacZ* (MBP151F/MBP193R primers), *PBAD-asdI-lacZ* (MBP151F/MBP151R primers) and *PBAD-asdII-lacZ*(MBP193F/MBP193R primers) were constructed by PCR amplification of desired fragments using primers containing homologies to P_BAD_ and *lacZ*. Similarly, transcriptional fusions P_BAD_-*asdI-II-lacZ* (MBP151F/MBP206R3 primers) and P_BAD_-*asdI-lacZ* (MBP151F/MBP206R1 primers) were generated by PCR amplification using a forward primer with homology to P_BAD_ and reverse primers containing *lacZ* RBS and *lacZ* homology. PCR products were then recombined into PM1205 strain using λ Red homologous recombination.

A plasmid harboring SgrS under the control of P*_teto-1_* promoter was constructed by PCR amplification of *sgrS* from *E. coli* MG1655 chromosomal DNA using primers containing NdeI and BamHI restriction sites. The resulting PCR product and vector pZA31R (23) were digested with NdeI and BamHI (New England Biolabs) restriction endonucleases. Digestion products were ligated using DNA Ligase (New England Biolabs) to produce plasmid pZAMB1 containing P_tet0-1_-sgrS allele.

Plasmid pZEMB8 containing P_lac0-1_-ptsG-gfp was constructed by PCR amplification of *ptsG* from MG1655 chromosomal DNA using primers containing KpnI and EcoRI restriction sites. The resulting PCR products and vector pZE12S (23) were digested with KpnI and EcoRI restriction endonucleases. Digestion products were ligated using DNA Ligase to produce pZEMB2. Superfolder *gfp* (referred to as *gfp*) was amplified from pXG10-SF (38) using primrs containing KpnI and XbaI restriction sites. pZEMB2 and the resulting PCR product were digested with KpnI and XbaI, and ligated with DNA ligase to produce pZEMB8. Plasmids with translational reporter fusions P_lac0-1_-*manX-gfp* (pZEMB10), P_lac0-1_-*yigL-gfp* (pZEMB15), P_lac0-1_-*purR-gfp* (pZEMB25), P_lac0-1_-*asdI-gfp* (pZEMB26) and P_lac0-1_-*asdI-II-gfp* (pZEMB27) were constructed by restriction cloning into pZEMB8 using KpnI and EcoRI restriction endonucleases.

### Media and reagents

Bacteria were cultured in Luria-Bertani (LB) broth medium or LB agar plates (35) at 37°C, unless stated otherwise. Bacteria were grown in MOPS (morpholine-propanesulfonic acid) rich defined medium (Teknova) with 0.2% fructose for reporter fluorescence assays. When necessary, media were supplemented with antibiotics at the following concentrations: 100 μg/ml ampicillin (Amp), 25 μg/ml chloramphenicol (Cm), 25 μg/ml kanamycin (Kan) and 50 μg/ml spectinomycin (Spec). Isopropyl β-D-1-thiogalactopyranoside (IPTG) was used at concentrations of 0.1-1.5 mM for induction of P*_lac0-1_* promoter, anhydrotetracycline (aTc) was used at 0-50 ng/ml for induction of P*_tet0-1_* promoter and L-arabinose was used at 0.000002%-0.2% for induction of P_BAD_ promoter, unless otherwise noted. To induce glucose-phosphate stress, 0.5% a-methylglucoside (αMG) was added to the growth medium.

### Reporter fluorescence assay

Bacterial strains were cultured overnight in MOPS rich medium supplemented with 0.2% fructose, Amp, and Cm, and diluted 1:100 into fresh medium with appropriate inducers (IPTG, aTc) in 48 well plates. Relative fluorescence units (RFU) and optical density (OD_600_) were measured over time. “GFP expression” was calculated by plotting RFU over OD_600_ and determining the slopes of linear regression plots for each IPTG concentration in exponentially growing cells in the presence of aTc to induce SgrS expression. “Promoter activity” was calculated by plotting RFU over OD_600_ and determining the slopes of linear regression plots for each IPTG concentration in exponentially growing cells in the absence of aTc.

### *In vitro* transcription and radiolabeling

Template DNA for *in vitro* transcription was generated by PCR using gene-specific oligonucleotides containing the T7 RNA polymerase promoter sequence. The following oligonucleotides were used to generate specific template DNA: MBP84F/MBP213R-ptsG (+1 to +240), O-JH218/MBP214R-*manX* (+1 to +240), MBP56F/MBP215R–*asdI-II* (+1 to +240), MBP56F/MBP222R–*asdI* (+1 to +110), MBP226F/MBP226R–*asdII* (+71 to +310), MBP65F/MBP174R–*purR* (+1 to +230), MBP216F/MBP216R–*yigL* (−191 to +50 relative to ATG translation start of *yigL*)MBP234F/MBP234R–*gfp* (+1 to +240) and O-JH219/O-JH119 were used to generate full-length *sgrS* template DNA. *In vitro* transcription of DNA templates was performed according to specifications of the MEGAscript T7 Kit (Ambion). *In vitro* transcribed RNA was 5’-end labeled with radioisotope ^32^P using the KinaseMax Kit (Ambion), according to the manufecturer’s instructions. Samples were purified by passing through Illustra ProbeQuant G-50 Micro Columns (GE Healthcare) followed by extraction with phenol-chloroform:isoamyl alcohol (Ambion), and labeled RNA was precipitated with ethanol:3M NaAc (30:1).

### RNA-RNA gel electrophoretic mobility shift assay

Different concentrations of unlabeled mRNA were mixed with 0.02 pmol of 5’-end labeled SgrS. Samples were denatured at 95°C for 1 min, placed on ice for 5 min, and incubated at 37°C for 30 min in 1x binding buffer (20 mM Tris-HCL [pH 8.0], 1mM DTT, 1 mM MgCl2, 20 mM KCl, 10 mM Na2HPO4 [pH 8.0]) (39). Non-denaturing loading buffer was added and samples were resolved for 6 h at 40 V on native 5.6% PAGE.

Protein-RNA gel electrophoretic mobility shift assay. 0.02 pmol of 5’-end labeled mRNA was denatured at 95°C for 1 min, and placed on ice for 5 min. Different concentrations of purified Hfq protein (His-tagged) were added. Samples were incubated at 37°C for 30 min in 1x binding buffer. Non-denaturing loading buffer was added and samples were resolved for 90 min at 20 mA on native 4.0% PAGE (39). SHAPE. The *asdI-II* RNA (0.15 μM) and SgrS RNA (0.075 μM, 0.15 μM, 0.30 μM, 0.75 μM, 1.5 μM, or 3.0 μM) were folded separately as in (40) using a modified SHAPE buffer (100 mM HEPES [pH 8.0], 2 mM MgCl2, 40 mM NaCl). For each SgrS concentration, the SgrS RNA or the equivalent volume of 0.5X TE was added to the *asdI-II* RNA and the samples were incubated at 37°C for 30 min. The RNAs were modified with *N*-methylisatoic anhydride (NMIA, 6.5 mM; Sigma-Aldrich) and collected by ethanol precipitation as in (40). Parallel primer extension inhibition and sequencing reactions were performed using fluorescently labeled primers complementary to the 3’ end of the *asdI-II* RNA (5′-AGATCAAAGGCATCCTGAAG, 22.5 nM; Applied Biosystems, ThermoFisher Scientific) as in (41) with minor modifications. Prior to primer binding, the RNAs were denatured and snap cooled and the reactions were carried out for 20 min at 52°C, followed by 5 min at 65°C. The cDNAs were analyzed on a 3730 DNA Analyzer (Applied Biosystems, Inc.). The data were processed and SHAPE reactivity (difference between the frequency of primer extension products at each nucleotide in +NMIA vs. −NMIA samples) was derived using the QuShape software (42). Data for each nucleotide were averaged with statistical outliers removed and normalized using the 2-8% rule (43). Relative reactivity was calculated by subtracting normalized SHAPE reactivity in the absence of the SgrS RNA from reactivity in the presence of the WT or MT SgrS RNA.

### Single-molecule fluorescence *in situ* hybridization (smFISH)

The *asdI-lacZ* (MB170), *asdII-lacZ* (MB183) and *asdI-II-lacZ* (MB171) strains were grown overnight at 37°C with shaking at 250 rpm in LB Broth Miller (EMD) containing Kan and Spec. The next day, the overnight cultures were diluted 100-fold into MOPS EZ rich defined medium (Teknova) with 0.2% (w/w) sodium succinate, 0.02% glycerol and L-arabinose (0.01% for the *asdI-lacZ* and *asdII-lacZ* strains, 0.002% for *asdI-II-lacZ*) and were allowed to grow at 37°C till the OD_600_ reached 0.15-0.25. αMG was added to the culture to a desired concentration to introduce sugar phosphate stress and induce SgrS sRNA expression. After 10 min of induction, the cells were fixed by mixing with formaldehyde (Fisher Scientific) at a final concentration of 4%.

Δ*sgrS* and Δ*lacZ* strains were grown in LB Broth Miller (EMD) at 37°C with shaking at 250 rpm overnight. The cultures were diluted 100-fold into MOPS EZ rich defined medium (Teknova) with 0.2% glucose and allowed to grow at 37°C until the OD_600_ reached 0.2. The cells were then fixed by mixing with formaldehyde at a final concentration of 4%. TK310 cells were grown overnight, similar to the knockout strains. The overnight culture was then diluted 100-fold into MOPS EZ rich defined medium (Teknova) with 0.2% glucose and 1 mM isopropyl β-D-1-thiogalactopyranoside (IPTG, Sigma-Aldrich) and allowed to grow at 37°C for 30 minutes. The cells were then fixed as described above.

The fixation and permeabilization of the cells were done using the methods published previously (44). After fixing with 4% formaldehyde, the cells were incubated at room temperature for 30 min. The cells were then centrifuged at 600 x g for 7 min and the pellets were washed with three times with 1X PBS 3. The cells were then permeabilized with 70% ethanol for 1 h at room temperature and stored at 4°C before fluorescence *in situ* hybridization.

The smFISH probes were designed using Stellaris Probe Designer and purchased from Biosearch Technologies (https://www.biosearchtech.com/). The labeling of the probes was performed using equal volumes of each probe. The final volume of sodium bicarbonate was adjusted to 0.1 M by adding 1/9 reaction volume of 1 M sodium bicarbonate (pH = 8.5). The probe solution was mixed with 0.05-0.25 mg of Alexa Fluor 647 or Alexa Fluor 568 succinimidyl ester (Life Technologies) dissolved in 5 μL DMSO. The dye was kept at 20-25 fold in molar excess relative to the probes. After incubation with gentle vortexing in the dark at 37°C overnight, the reaction was quenched by adding 1/9 reaction volume of 3 M sodium acetate (pH 5). Unconjugated dye was removed by ethanol precipitation followed by P-6 Micro Bio-Spin Column (Bio-Rad).

A previously published protocol (44) was used for the hybridization procedure. 60 μ of permeabilized cells were washed with FISH wash solution (10% formamide in 2X SSC [Saline Sodium Citrate] buffer) and resuspended in 15 μ hybridization buffer (10% dextran sulfate, 1 mg/ml *E. coli* tRNA, 0.2 mg/ml BSA, 2 mM vanadyl ribonucleoside complexes, 10% formamide in 2X SSC) with probes. Nine probes labeled with Alexa Fluor 647 were used for sRNA SgrS and 24 probes labeled with Alexa Fluor 568 were used for mRNA *lacZ*. The concentration of the labeled probes for SgrS and *lacZ* mRNA were 50 nM and 15 nM, respectively. The reactions were incubated in the dark at 3 °C overnight. The cells were then resuspended in 20X volume FISH wash solution and centrifuged. They were then resuspended in FISH wash solution, incubated for 30 min at 30°C and centrifuged and this was repeated 3 times. The cells were pelleted after the final washing step and resuspended in 20 μl 4X SSC and stored at 4°C for imaging. The labeled cells were immobilized on a poly-L-lysine (Sigma-Aldrich) treated 1.0 borosilicate chambered coverglass (Thermo Scientific™ Nunc^TM^ Lab-Tek^TM^). They were then imaged in imaging buffer (50 mM Tris-HCl [pH = 8.0], 10% glucose, 1% β-mercaptoethanol [Sigma-Aldrich], 0.5 mg/ml glucose oxidase [Sigma-Aldrich] and 0.2% catalase [Calbiochem] in 2X SSC).

### Single-molecule localization-based super-resolution imaging

An Olympus IX-71 inverted microscope with a 100X NA 1.4 SaPo oil immersion objective was used for the 3D super-resolution imaging. The lasers used for two-color imaging were Sapphire 568–100 CW CDRH, Coherent (568nm) and DL640-100-AL-O, Crystalaser (647nm) and DL405-025. Crystalaser (405nm) was used for the reactivation of Alexa 647 and Alexa 568 fluorophores. The laser excitation was controlled using mechanical shutters (LS6T2, Uniblitz). A dichroic mirror (Di01-R405/488/561/635, Semrock) was used to reflect the laser lines to the objective. The objective collected the emission signals and then they made their way through an emission filter (FF01-594/730-25, Semrock for Alexa 647 or HQ585/70M 63061, Chroma for Alexa 568) and excitation laser was cleaned up using notch filters (ZET647NF, Chroma, NF01-568/647-25x5.0 and NF01-568U-25, Semrock). Samples were then imaged on a 512x512 Andor EMCCD camera (DV887ECS-BV, Andor Tech). Astigmatism was introduced by placing a cylindrical lens with a focal length of 2 m (SCX-50.8-1000.0-UV-SLMF-520-820, CVI Melles Griot) in the emission path between two relay lenses with focal lengths of 100 mm and 150 mm each, which facilitated 3D imaging. In this setup, each pixel corresponded to 100 nm. We used the CRISP (Continuous Reflective Interface Sample Placement) system (ASI) to keep the z-drift of the setup to a minimum. The image acquisition was controlled using the storm-control software written in Python by Zhuang and coworkers (45, 46) and available at GitHub.

The imaging of the sample began with a DIC image of the sample area. Subsequently two-color super-resolution imaging was performed. 647nm excitation was used first and after image acquisition was completed for Alexa Fluor 647, 568nm excitation was used to image Alexa Fluor 568. 405 nm laser power was increased slowly to compensate for fluorophore bleaching and also to maintain moderate signal density. Imaging was stopped when most of the fluorophores had photobleached and the highest reactivation laser power was reached.

The raw data acquired using the acquisition software were analyzed using the method described in previously published work (29), which was a modification of the algorithm published by Zhuang and coworkers (45, 46). The clustering analysis on the localization data was performed using MATLAB codes as described previously (29). Background signal was estimated using Δ*sgrS* and Δ*lacZ* strains and they were prepared, imaged and analyzed as described above. TK310 cells were prepared, imaged and analyzed in the same way as a low copy *lacZ* mRNA sample for copy number calculation. The copy number calculation was also performed using MATLAB codes as described previously (29).

## ACKNOWLEDGEMENTS

We would like to extend a special thank you to Erel Levine for providing plasmids. We are grateful to Jennifer Rice, Rich Yemm, Divya Balasubramanian, Chelsea Lloyd, Alisa King, Jessica Kelliher and other current and past members of the Vanderpool lab for strains, plasmids and valuable advice. We appreciate and thank Prof. James Slauch and members of his lab for fruitful discussions.

